# The direct interaction with transcriptional factor TEAD4 implied a straightforward regulation mechanism of tumor suppressor NF2

**DOI:** 10.1101/2022.11.28.518166

**Authors:** Liqiao Hu, Mengying Wu, Lingli He, Liang Yuan, Lingling Yang, Bin Zhao, Lei Zhang, Xiaojing He

**Author notes:** Correspondence: Xiaojing He. These authors contributed equally (Liqiao Hu, Mengying Wu, Lingli He).

## Abstract

As an output effecter of Hippo signaling pathway, the transcription factor TEAD and co-activator YAP play crucial functions in promoting cell proliferation and organ size. The tumor suppressor NF2 has been shown to activate LATS1/2 kinases and interplay with Hippo pathway to suppress YAP-TEAD complex. But, whether and how NF2 could directly regulate TEAD remains unknown. We identified a direct link and physical interaction between NF2 and TEAD4. NF2 interacted with TEAD4 through its FERM domain and the C-terminal tail, and decreased protein stability of TEAD4 independently of LATS1/2 and YAP. Furthermore, NF2 inhibited TEAD4 palmitoylation and retained the cytoplasmic translocation of TEAD4, resulting in ubiquitination and dysfunction of TEAD4. Moreover, the interaction with TEAD4 is required for NF2 function to suppress cell proliferation. These findings revealed a new role of NF2 as a binding partner and inhibitor of the transcription factor TEAD, and would shed light on an alternative mechanism of how NF2 functions as a tumor suppressor through the Hippo signaling cascade.

## Introduction

In multicellular animals, cell proliferation and death must be precisely coordinated to ensure proper organ size and tissue homeostasis. The Hippo signaling pathway was initially identified as a key determinant of organ size (Harvey et al., 2003; Huang et al., 2005; Pan, 2010; Wu et al., 2003). This pathway is highly conserved from *Drosophila* to mammals (Yu et al., 2015; Zhao et al., 2010a). The Hippo pathway constitutes a major kinase cascade, including the mammalian STE20-like protein kinase 1/2 (MST1/2) and large tumor suppressor kinase1/2 (LATS1/2), which inhibit two transcriptional co-activators, Yes-associated protein (YAP) and transcriptional co-activator with PDZ-binding motif (TAZ), via phosphorylation (Zhao et al., 2010b). Dephosphorylated and activated YAP/TAZ translocate into the nucleus, where they interact with the TEA domain transcription factors (TEADs), and induce the expression of target genes, such as *CTGF* and *CYR61*, to modulate cell proliferation, differentiation, and tumorigenesis (Ota and Sasaki, 2008; Zhang et al., 2008; Zhao et al., 2008). Unlike *Drosophila*, which expresses only one TEAD homolog, Scalloped (Sd), there are four TEAD homologs in mammals (TEAD1, TEAD2, TEAD3, and TEAD4). TEADs share a similar domain structure: a DNA-binding domain (DBD) at the N-terminus and a YAP-binding domain (YBD) at the C-terminus (Huh et al., 2019; Zhang et al., 2008). Dysregulation of the Hippo pathway has been linked to many human diseases, and targeted inhibition of the YAP-TEAD transcriptional complex for cancer therapy is being actively explored (Dey et al., 2020; Yu et al., 2015; Zheng and Pan, 2019).

In addition to YAP/TAZ, the transcriptional activity of TEADs is regulated by different binding factors, including VGLL4, glucocorticoid receptor(GR), TCF4, and AP-1(He et al., 2019; Jiao et al., 2017, 2014; Liu et al., 2016). Specifically, VGLL4 directly competes with YAP/TAZ for binding to TEADs, thereby suppressing their transcriptional activity (Deng and Fang, 2018). P38 binding-dependent cytoplasmic translocation of TEADs provides spatial modulation of transcriptional activity (Lin et al., 2017). Post-translational modifications of TEADs, such as phosphorylation and palmitoylation, govern their protein stability and activity (Chan et al., 2016; Gupta et al., 2000; Jiang et al., 2001; Noland et al., 2016). Four TEAD homologs have been found to be palmitoylated in mammalian cells (Kim and Gumbiner, 2019; Mesrouze et al., 2017), and palmitoylation of TEAD is critical for protein stability and YAP-TEAD interaction (Noland et al. 2016; Chan et al. 2016). Although targeting TEAD palmitoylation is considered as a potential strategy for Hippo pathway molecular therapy (Bum-Erdene et al., 2019; Pobbati et al., 2015), the mechanisms regulating TEAD palmitoylation and depalmitoylation remain unclear.

Neurofibromin 2 (NF2), also called Merlin, is an **E**zrin, **R**adixin, and **M**oesin(ERM) family protein that acts as a tumor suppressor, and the development of schwannoma, meningioma, ependymoma, and malignant mesothelioma in humans is highly associated with loss-function and mutations of *NF2* (Chen et al., 2017; Cheng et al., 1999; Kalamarides, 2002). NF2 functions in the Hippo pathway by responding to extracellular stimuli, such as cell density and osmotic stress (Cooper and Giancotti, 2014; Hong et al., 2020). NF2 associates with LATS1/2, thus activating the major kinase cascade of Hippo pathway to inhibit YAP/TAZ and suppress cell proliferation and tumorgenesis (Yin et al., 2013). Several binding partners of NF2, including Angiomotin (AMOT) and E3 ubiquitin ligase CRL4^DCAF1^, are also involved in modulating Hippo pathway (Li et al., 2010, 2015). However, whether NF2 directly regulates TEADs remains unclear and how NF2 modulates the Hippo pathway is not yet fully understood.

In this study, we identified the physical interaction between tumor suppressor NF2 and transcription factor TEAD4. We found that NF2 directly interacted with TEAD4 through its FERM domain and the C-terminal tail, and decreased protein stability of TEAD4 independently of LATS1/2 and YAP. We further revealed the molecular mechanism that NF2 inhibited TEAD4 palmitoylation and retained its cytoplasmic translocation via the direct interaction, resulting in ubiquitination and dysfunction of TEAD4. Moreover, the TEAD4 interaction is required for NF2 function to suppress tumor cell proliferation. These findings implied a new role of NF2 as a binding partner and inhibitor of TEADs, and expanded the molecular mechanism of how NF2 functions as a tumor suppressor.

## Results

### NF2 decreases the protein levels of TEADs independently of LATS1/2 and YAP

As an upstream activator in the Hippo signaling pathway, the tumor suppressor NF2 has been shown to activate LATS1/2 kinases and suppress YAP function (Yin et al., 2013). NF2 also interacts with other regulators, AMOT and DCAF1, to modulate the Hippo pathway (Li et al., 2010, 2015). We were curious whether other effectors might be directly involved in NF2 function. Firstly, we examined the protein levels of major Hippo pathway components in HEK293T cells with NF2 overexpression. Consistent with a previous report (Yin et al., 2013), NF2 promoted YAP phosphorylation (**Figure 1A**). Unexpectedly, the protein levels of TEAD4 were markedly reduced along with NF2 overexpression (**Figure 1A**). We then confirmed this result by silence and rescue experiments. Depletion of *NF2* by small interfering RNA (siRNA) induced an increase in both TEAD2 and TEAD4 protein levels, and the re-expression of NF2 decreased their protein levels again (**Figure supplement 1A**). Notably, the mRNA levels of TEADs were unaffected by NF2 overexpression (**Figure supplement 1B**). These results suggest a hypothesis that NF2 may regulate protein levels of TEADs instead of transcription levels.

**Figure 1.**
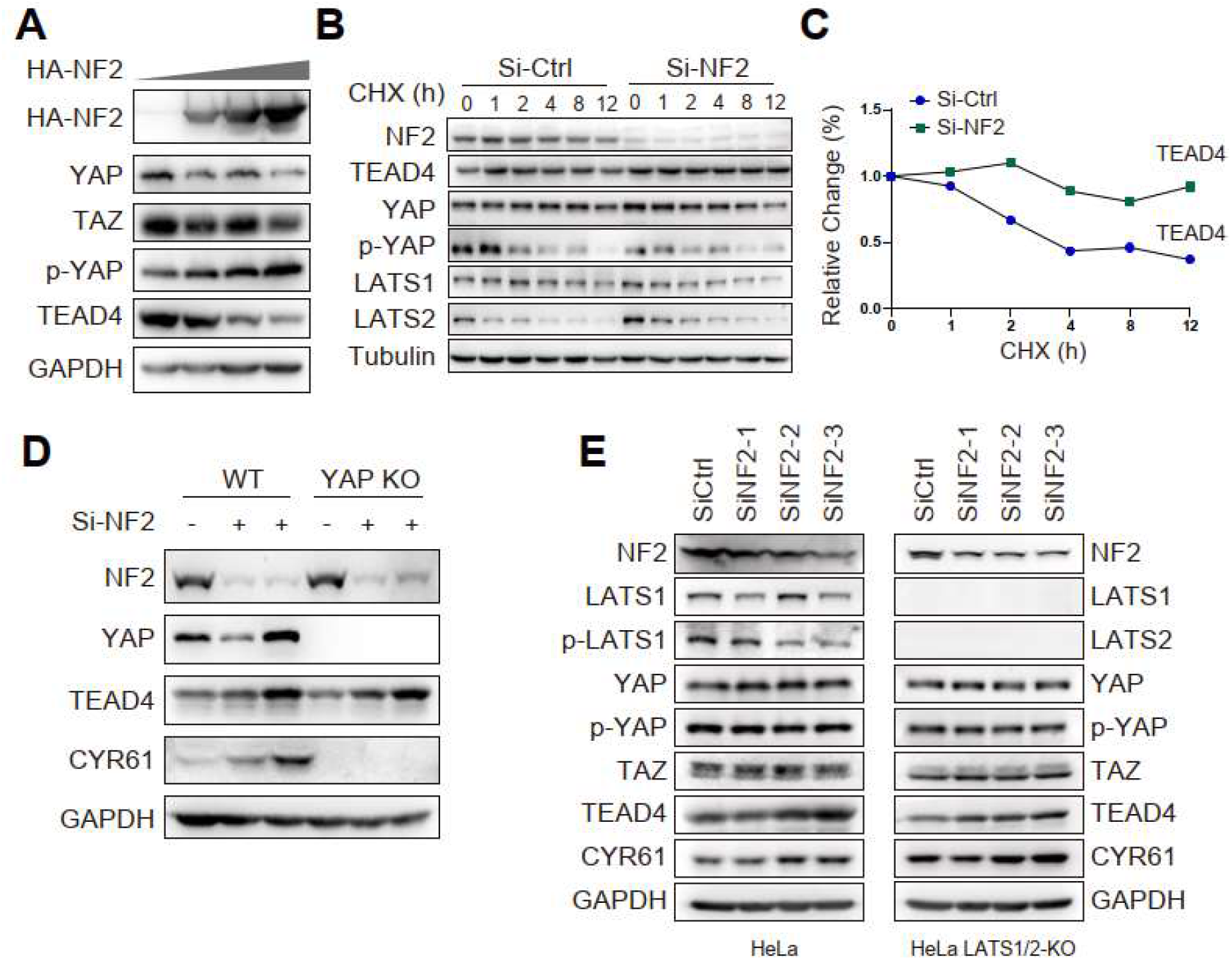
NF2 decreases the protein levels of TEADs independently of LATS1/2 and YAP. (A) Protein levels of TEAD4, YAP and TAZ were determined via western blotting in HEK293T cells with overexpression of HA-NF2. (B) MCF-10A cells were transfected with the control siRNA or NF2 siRNA. Then, 100 ug/mL of cycloheximide (CHX) was added and the cells were harvested at the indicated time points. Protein levels of endogenous TEAD4, YAP, p-YAP, and LATS-1/2 were determined via western blotting. (C) Quantitative analysis of TEAD4 protein level in (B). (D) Wild-type (WT) and *YAP* knockout (KO) HeLa cells were transfected with or without the NF2 siRNA. Protein levels of TEAD4, YAP, and CYR61 were determined via western blotting. (E) WT and *LATS*1/2 KO HeLa cells were transfected with the control siRNA or NF2 siRNA. Protein levels of TEAD4, YAP/TAZ, LATS1, and CYR61 were determined by western blotting. **Figure 1-source data 1**. Whole uncropped blots represented in ***Figure 1A***. NF2, YAP, TAZ, p-YAP, TEAD4 and GAPDH protein levels in HEK293T cells. **Figure 1-source data 2**. Whole uncropped blots represented in ***Figure 1B***. NF2, YAP, p-YAP, TEAD4, Lats1, Lats2 and Tubulin protein levels in MCF-10A cells. **Figure 1-source data 3**. Whole uncropped blots represented in ***Figure 1D***. NF2, TEAD4, YAP, CYR61 and GADPH protein levels in Wild-type (WT) and *YAP* knockout (KO) HeLa cells. **Figure 1-source data 4**. Whole uncropped blots represented in ***Figure 1E***. NF2, Lats1, Lats2, p-Lats1, TEAD4, YAP, p-YAP, TAZ, CYR61 and GADPH protein levels in Wild-type (WT) and *LATS*1/2 knockout (KO) HeLa cells.

We then examined the protein stability of TEAD4 in the cells treated with cycloheximide (CHX) to inhibit *de novo* protein synthesis. In comparison with control cells, knockdown of *NF2* significantly increased the half-life of TEAD4 protein (**Figure 1B and 1C**). The protein levels of other components in Hippo pathway, YAP and Lats1/2, were also detected, and exhibited similar levels in both control and NF2 knockdown cells. Thus, it suggests that NF2 decreased TEAD4 protein level by altering the protein stability of TEAD4.

Since YAP is a well-known major partner of TEAD4, we then asked if YAP is involved in the regulation of TEAD4 protein stability by NF2. We generated *YAP* knockout (KO) HeLa cells by CRISPR/Cas9, in which successful *YAP* KO were verified by western blotting (**Figure 1D**). The knockdown of *NF2* robustly increased TEAD4 protein levels in both WT and *YAP* KO cells (**Figure 1D**), and NF2 overexpression also decreased TEAD4 protein levels in *YAP* KO cells (**Figure supplement 1C**), indicating that NF2 decreased the protein levels of TEADs independently of YAP. We also wondered if LATS1/2 might be involved in the regulation of TEAD4 protein level by NF2. Again, in both control and *LATS1/2* KO cells, knockdown of *NF2* increased the TEAD4 protein expression levels (**Figure 1E**). Taken together, our results suggest a direct regulation that NF2 could decrease the protein level and stability of TEAD4 independently of LATS1/2 and YAP.

### NF2 physically interacts with TEAD4 through its FERM domain and C-terminal tail

To further explore the direct link between NF2 and TEAD4, we purified the YBD domain of TEAD4 (TEAD4-YBD) with SUMO-His tag and full-length NF2 with GST tag from *Escherichia coli*, and examined their interaction using a glutathione S-transferase (GST) pull-down assay. Indeed, TEAD4-YBD strongly bound to GST-NF2 *in vitro* (**Figure 2A**), highly suggesting a direct and physical interaction between NF2 and TEAD4.

**Figure 2.**
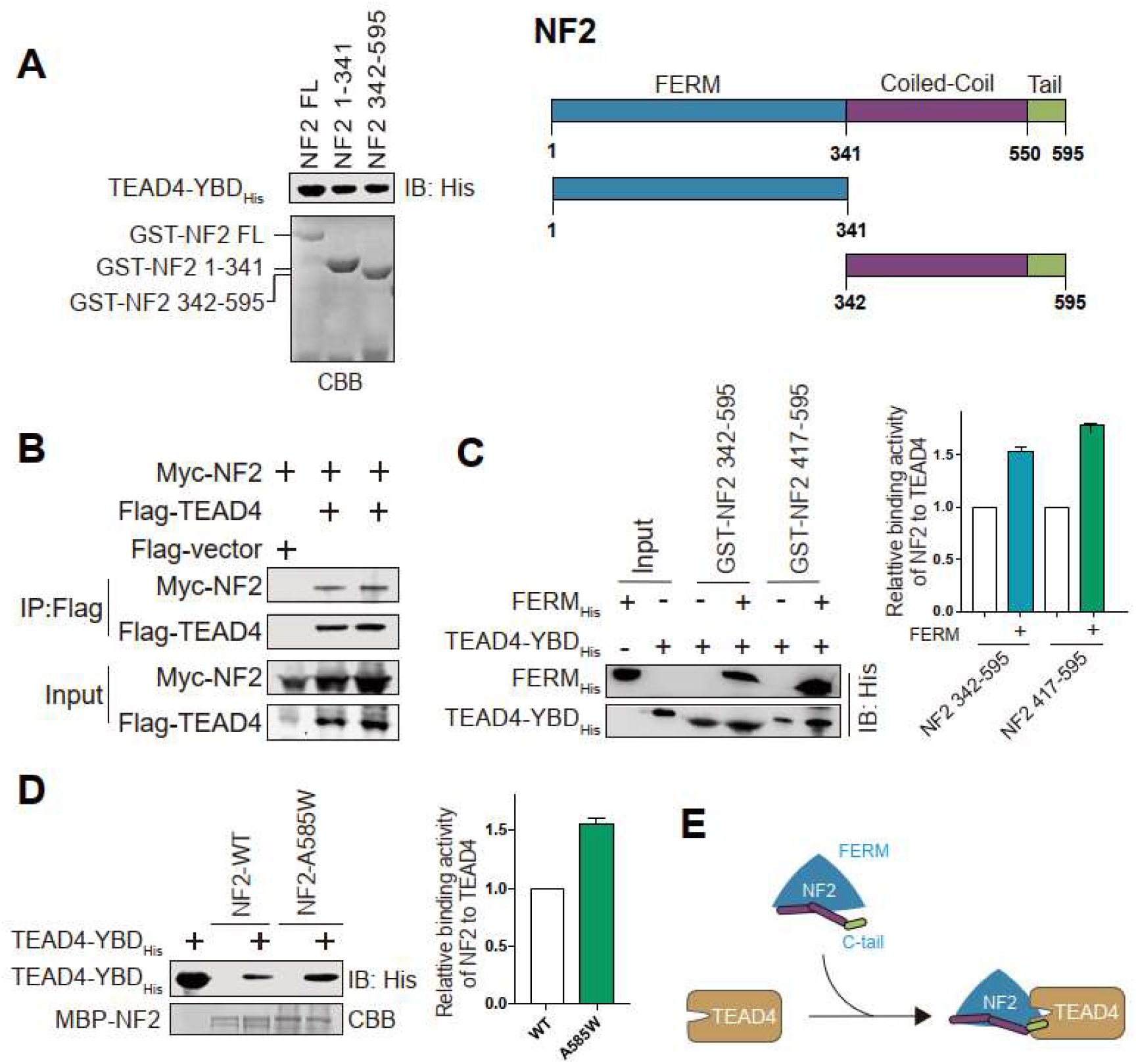
NF2 interacts with TEAD4 through FERM domain and C-terminal tail. (A) The GST-pull down assay was performed to assess the interaction between SUMO-tagged TEAD4-YBD and GST-tagged NF2 truncations, and schematic views of NF2 full-length and truncations showed in right panel. CBB, Coomassie brilliant blue. (B) Co-immunoprecipitation experiment of Myc-tagged NF2 with Flag-tagged TEAD4 was performed in HEK293T cells. Cell lysates were treated to anti-Flag beads and immunoblotted with indicated antibodies. Expression of Flag vector was set as control. (C) *In vitro* binding assay of GST-tagged NF2 C-terminal fragments with His-TEAD4-YBD, the FERM domain enhanced the interaction between TEAD4 and NF2 C-terminal fragments. Quantitative analysis of relative protein binding activity of NF2 to TEAD4 was shown in the right panel. Mean ± s.e.m, N = 3. (D) *In vitro* pull-down assay of MBP-NF2 WT and A585W mutant with His-TEAD4-YBD to assess the interaction between TEAD4 and NF2. CBB, Coomassie brilliant blue. Quantitative analysis of relative protein binding activity of NF2 to TEAD4 was shown in the right panel. Mean ± s.e.m, N = 3. (E) A cartoon model of NF2 binding to TEAD4 through FERM domain and C-terminal tail. **Figure 2-source data 1**. Whole SDS-PAGE images and uncropped blots represented in ***Figure 2A***. TEAD4-YBD_His_, GST-NF2-FL, GST-NF2 1-341, GST-NF2 342-595 protein levels in GST pull-down assay. **Figure 2-source data 2**. Whole uncropped blots represented in ***Figure 2B***. Flag-TEAD4 and Myc-NF2 protein levels with Co-immunoprecipitation assay in HEK293T cells. **Figure 2-source data 3**. Whole uncropped blots represented in ***Figure 2C***. TEAD4-YBD_His_, NF2 FERM_His_ protein levels in GST pull-down assay. **Figure 2-source data 4**. Whole SDS-PAGE images and uncropped blots represented in ***Figure 2D***. TEAD4-YBD_His_ and MBP-NF2 protein levels in MBP pull-down assay.

According to the domain organization of NF2 protein which containing the N-terminal FERM domain, central coiled-coil domain, and C-terminal tail(Li et al., 2015), seven truncations were constructed to test their interaction with TEAD4-YBD (**Figure 2A and supplement 2A**). In the GST pull-down assay, both the N-terminal FERM domain (1-341 aa) and C-terminal half (342-595 aa) remained to interact with TEAD4-YBD at similar level *in vitro* (**Figure 2A**). Among the C-terminal half, the tail region (C-tail, 550-595 aa) still bound to TEAD4 (**Figure supplement 2A**). The interaction between NF2 and TEAD4 full length was verified by co-immunoprecipitation assay in HEK293T cells (**Figure 2B**), which further confirmed the direct and physical interaction between NF2 and TEAD4.

Intramolecular interaction of NF2 has been suggested to be formed by FERM domain and C-terminal tail (Chinthalapudi et al., 2018; Li et al., 2015; Sher et al., 2012). We then wondered if the intramolecular interaction of NF2 might affect its interaction with TEAD4. In comparison with C-terminal fragments alone, co-incubation of FERM domain markedly enhanced their binding to TEAD4 (**Figure 2C**), indicating that the intramolecular interaction of NF2 increased the interaction with TEAD4. Moreover, the A585W mutation of NF2, which could stabilize the intramolecular interaction and be inactive for LATS1/2 interaction (Li et al., 2015), exhibited stronger binding capacity to TEAD4 than NF2-WT in MBP pull-down assay (**Figure 2D**). Taken together, these binding results suggest that NF2 directly interacts with TEAD4 through both FERM domain and the C-terminal tail (**Figure 2E**).

### Characterization of the interaction between NF2 and TEAD4

Given that NF2 is a novel binding partner of TEAD4, we next characterized the interaction interface of NF2. Based on the crystal structure of NF2(Li et al., 2015), single or combined point mutations on the structural surface of NF2 protein were designed for the binding screen (**Figure supplement 2B**). Collectively, the binding assay pinpointed that L297, I301, and H304 residues in the FERM domain F3 lobe, L582 and F591 residues in the C-tail of NF2 mediated its interaction with TEAD4 (**Figure 3A, 3B and supplement 2C**). We then generated two grouped mutants NF2-5A (L297A/I301A/H304A/L582A/F591A) and NF2-4A-del (L297A/I301A/H304A/L582A and deletion of 590-595 aa) (**Figure supplement 2C**), and both mutants abolished their ability to interact with TEAD4 in co-immunoprecipitation assay (**Figure 3C**). Thus, the key residues L297/I301/H304 on FERM domain and L582/F591 on C-tail were identified to mediate the interaction between NF2 and TEAD4.

**Figure 3.**
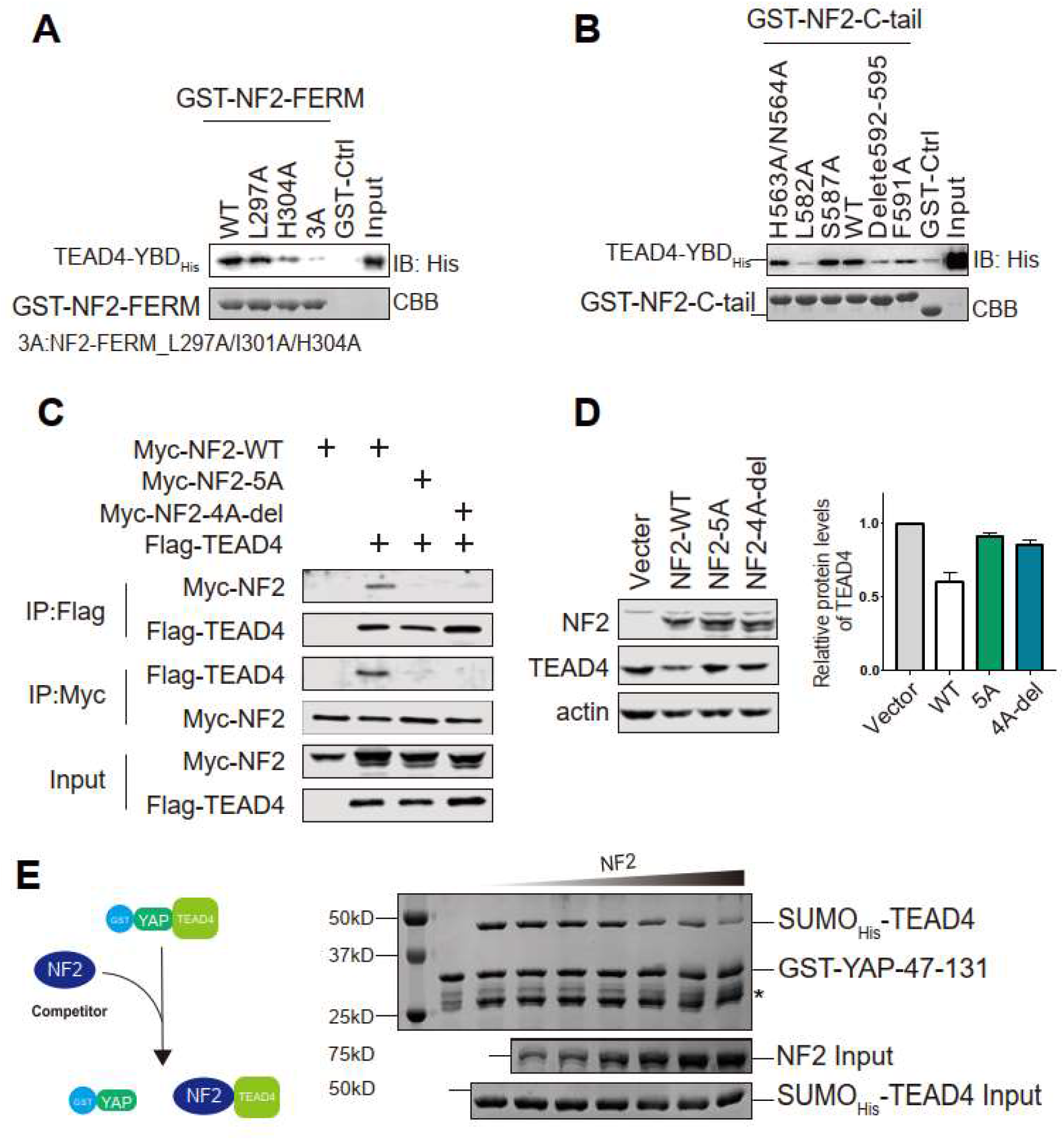
NF2 decreases TEAD4 protein level via direct interaction. (A) GST pull-down assay of GST-NF2-FERM WT and mutants with His-TEAD4-YBD was performed to assess the interaction between TEAD4 and NF2-FERM. (B) GST pull-down assay of GST-NF2-507-595 WT and mutants with His-TEAD4-YBD was performed to assess the interaction between TEAD4 and NF2 C-terminal tail. (C) Co-immunoprecipitation experiment of Myc-tagged NF2 WT and mutants with Flag-tagged TEAD4 was performed in HEK293T cells. Cell lysates were treated to anti-Flag or anti-Myc beads and immunoblotted with indicated antibodies. (D) TEAD4 Protein levels in NCI-H226 cells overexpressing Myc-tagged NF2 WT or mutants were determined via western blotting by indicated antibody. Quantitative analysis of relative protein levels of TEAD4 was shown in the right panel. Mean ± s.e.m, N = 3. (E) Competitive binding assay was performed to detect the binding effect of YAP-TEAD4 complex with a dose addition of NF2. **Figure 3. source data 1**. Whole SDS-PAGE images and uncropped blots represented in ***Figure 3A***. TEAD4-YBD_His_ and GST-NF2 FERM protein levels in GST pull-down assay. **Figure 3-source data 2**. Whole SDS-PAGE images and uncropped blots represented in ***Figure 3B***. TEAD4-YBD_His_ and GST-NF2 507-595 protein levels in GST pull-down assay. **Figure 3-source data 3**. Whole uncropped blots represented in ***Figure 3C***. Flag-TEAD4 and Myc-NF2 protein levels with Co-immunoprecipitation assay in HEK293T cells. **Figure 3-source data 4**. Whole uncropped blots represented in ***Figure 3D***. TEAD4, Myc-NF2 and beta-actin protein levels in NCI-H226 cells. **Figure 3-source data 5**. Whole SDS-PAGE images represented in ***Figure 3E***. TEAD4-YBD_His_, GST-YAP 47-131 and NF2 protein levels in Competitive binding assay.

We then asked if these binding-deficient mutants affect the function of NF2 to decreaseTEAD4 protein level. We introduced NCI-H226 cell line, an NF2-non-expressing malignant pleural mesothelioma (MPM) cell line, which could exclude the effect of endogenous NF2. In comparison with control cells, overexpression of NF2-WT decreased TEAD4 protein level to 60% (**Figure 3D**). Consistent with their interaction deficiency withTEAD4, NF2-5A and NF2-4A-del mutants restored TEAD4 protein level around 85% (**Figure 3D**), suggesting that NF2 decreased TEAD4 protein level via direct interaction.

Since both YAP and NF2 bind to the YBD domain of TEAD4, we explored whether NF2 and YAP bound to the same surface on TEAD4. The *in vitro* competitive binding assay was performed and showed that NF2 gradually competed off TEAD4 from GST-YAP in a dose-dependent manner (**Figure 3E**), indicating that NF2 and YAP occupied the same interface on TEAD4-YBD domain and NF2 potentially inhibited the formation of YAP-TEAD complex.

### NF2 induces the cytoplasmic retention of TEAD4 via interaction

TEAD4 functions as a transcription factor in the nucleus, while NF2 is a plasma membrane-associated protein (Yin et al., 2013). To characterize the spatial localization of the NF2-TEAD4 complex in cell, we performed bimolecular fluorescence complementation (BiFC) assays, in which two non-fluorescent half fragments of the yellow fluorescent protein (YFP) were fused with two binding partners, respectively. No fluorescence was detected upon co-expression of NF2-nYFP and TEAD4-nYFP in HEK293 cells, similar to control cells expressing NF2-cYFP or TEAD4-cYAP alone **(Figure 4A)**. Co-expression of NF2-cYFP and LATS2-nYFP, which are well-known binding partners, resulted in fluorescence at the plasma membrane **(Figure 4A)**. Co-expression of NF2-cYFP and TEAD4-nYFP also resulted in YFP signals at the plasma membrane, but more in the cytoplasm **(Figure 4A)**, suggesting that they form a complex in the cytoplasm rather than in the nucleus. Fluorescence signals were sequentially quantified by flow cytometry **(Figures 4B and supplement 3A)**, validating that NF2 strongly interacted with TEAD4 in cells, as NF2 and LATS2 did.

**Figure 4.**
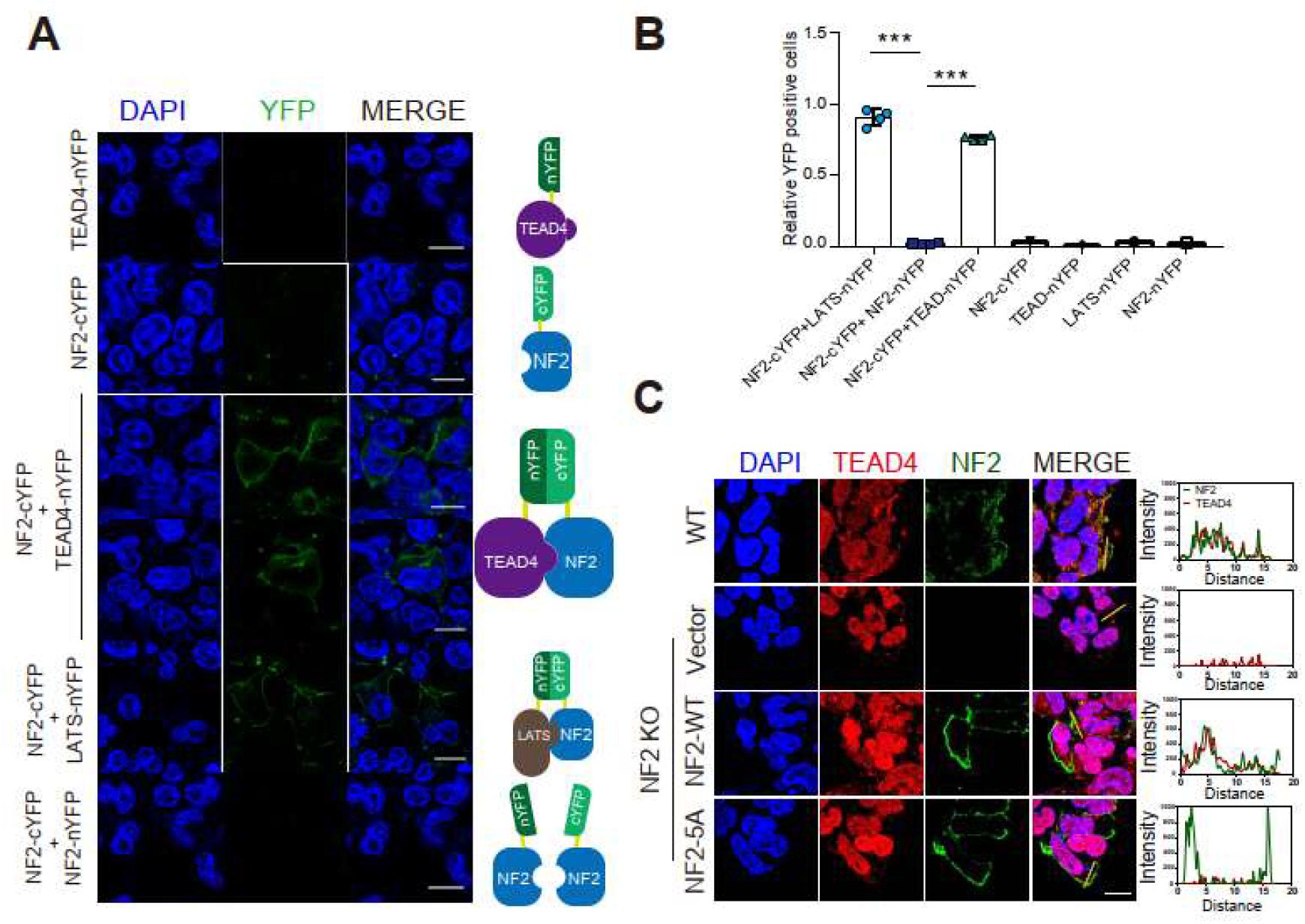
NF2 induces the cytoplasmic retention of TEAD4 via interaction. (A) BiFC assays were performed to detect the location of the NF2-TEAD4 complex. HEK293T cells were transfected with TEAD4-nYFP, NF2-cYFP, NF2-cYFP and TEAD4-nYFP, NF2-cYFP and Lats-nYFP, and NF2-cYFP and NF2-nYFP. The fluorescence signals of intact YFP were detected. Representative images are shown here. Scale bar = 10 μm. (B) Quantification of flow cytometry from BiFC assay (A). Mean ± s.e.m. \*\*\**P* < 0.001. (C) NF2 induces the cytoplasmic translocation of TEAD4. Immunofluorescence of endogenous TEAD4 was detected in *NF2*-KO HEK293A cells with expression of Myc-NF2 WT and mutant. Scale bar = 10μm. **Figure 4-source data 1**. Source data for quantifications graphed in ***Figure 4B***.

To examine whether NF2 induces the translocation of TEAD4 via the direct interaction, we detected the subcellular localization of TEAD4 by immune-fluorescence in the *NF2*-KO HEK293A cells with overexpression of NF2-WT and NF2-5A, respectively. Compared with the nuclear localization of TEAD4 in *NF2*-KO cells, clear fluorescence signals of TEAD4 were visible in the cytoplasm of NF2-WT expressed cells **(Figure 4C)**. However, the fluorescence signal of TEAD4 could not be detected in the cytoplasm of NF2-5A expressed cells **(Figure 4C)**, suggesting that NF2 induces the cytoplasmic retention of TEAD4 through the direct interaction.

### NF2 inhibits TEAD4 palmitoylation and presumably causes the sequential ubiquitination

As palmitoylation of TEAD4 is required for its protein stability (Chan et al., 2016; Kim and Gumbiner, 2019; Noland et al., 2016), we further investigated whether NF2 decreased protein stability of TEAD4 through palmitoylation. The *in vitro* auto-palmitoylation assays were performed by click chemistry-based methods (Zheng et al., 2015)(**Figure 5A**). TEAD4 auto-palmitoylation was significantly decreased along with NF2 incubation, but not YAP (**Figure 5B and 5C**), indicating that NF2 directly inhibited the auto-palmitoylation of TEAD4 *in vitro*.

**Figure 5.**
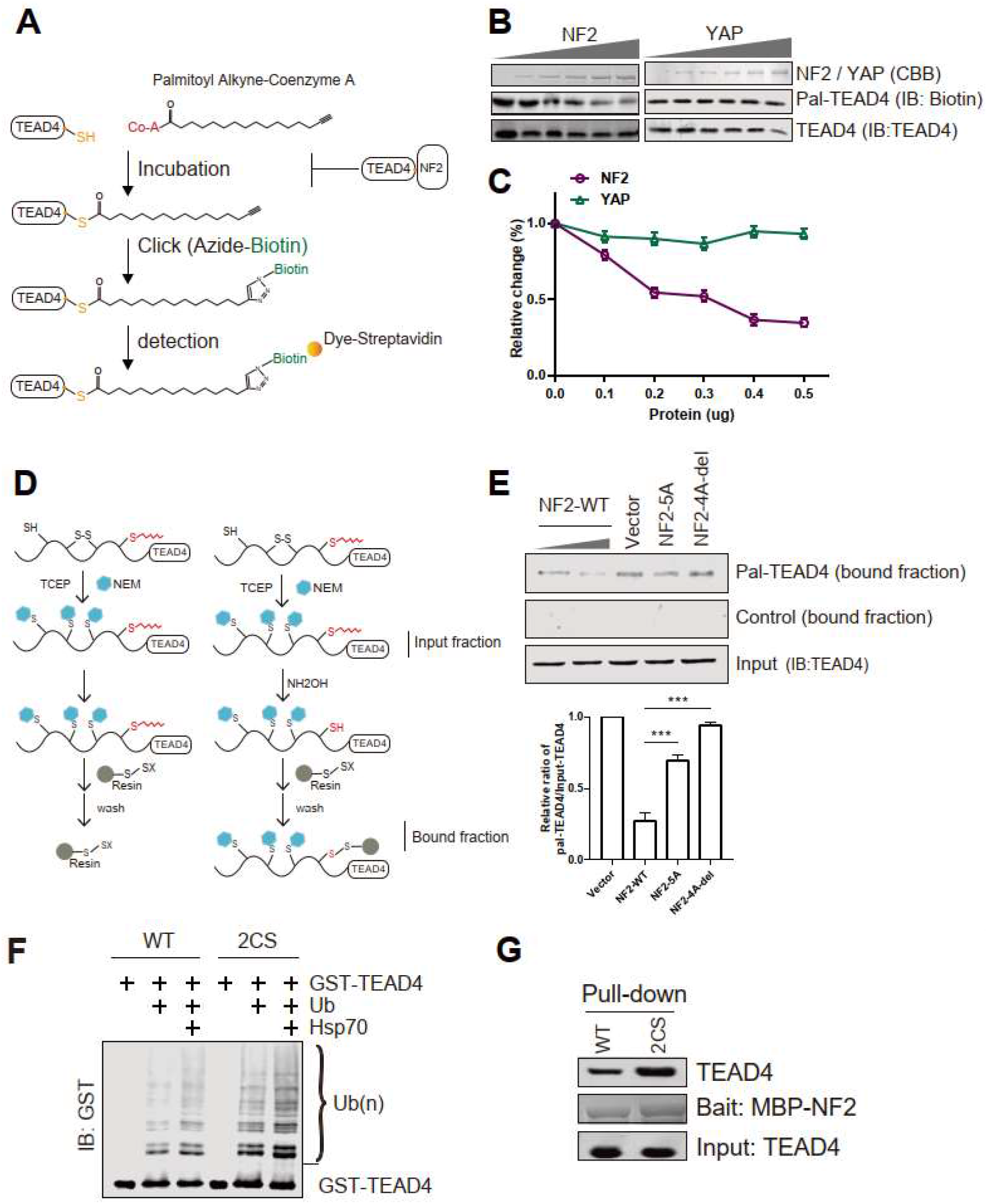
NF2 inhibits TEAD4 palmitoylation via direct interaction. (A) Schematic model of the *in vitro* auto-palmitoylation assay of recombinant TEAD4. (B) *In vitro* auto-palmitoylation assay of recombinant TEAD4 incubated with NF2 or YAP was performed under protocols. Palmitoylation levels were detected via streptavidin blotting. (C) Quantitative analysis of palmitoylated TEAD4 (Pal-TEAD4) panels in (B). (D) Schematic model of the *in vivo* palmitoylation assay with acyl resin-assisted capture methods in cells. (E) *In vivo* palmitoylation assay to detect the palmitoylation levels of endogenous TEAD4 in NCI-H226 cells expressing the Myc-NF2 WT and mutants. palmitoylation levels of TEAD4 were determined via western and streptavidin blotting. Quantitative analysis of palmitoylated TEAD4 (Pal-TEAD4) was shown in the down panel. Mean ± s.e.m, N = 3, \*\*\**P*< 0.001. (F) *In vitro* ubiquitination assay was performed with purified recombinant E1, UbcH5b as the E2 ubiquitin-conjugating enzyme, and E3 ligase CHIP to detect the ubiquitination level of TEAD4 WT and 2CS. (G) *In vitro* pull-down assay of MBP-NF2 withTEAD4 WT and 2CS. CBB, Coomassie brilliant blue. **Figure 5-source data 1**. Whole SDS-PAGE images and uncropped blots represented in ***Figure 5B***. Pal-TEAD4-YBD_His_, TEAD4-YBD_His_, MBP-NF2 FL and YAP FL protein levels in *In vitro* auto-palmitoylation assay. **Figure 5-source data 2**. Whole uncropped blots represented in ***Figure 5E***. Pal-TEAD4 and TEAD4 protein levels in NCI-H226 cells. **Figure 5-source data 3**. Whole uncropped blots represented in ***Figure 5F***. Ubiquitination levels of GST-TEAD4 in *In vitro* ubiquitination assay. **Figure 5-source data 4**. Whole SDS-PAGE images and uncropped blots represented in ***Figure 5G***. TEAD4-YBD_His_ and MBP-NF2 FL protein levels in MBP pull-down assay.

Next, we examined the palmitoylation of TEAD4 in NCI-H226 cells using an acyl resin-assisted capture assay (Forrester et al., 2011) (**Figure 5D**). Notably, the expression of NF2-WT dramatically reduced TEAD4palmitoylation, but NF2-5A and NF2-4A-del mutants effected TEAD4 palmitoylation slightly in cells (**Figure 5E**), implying that NF2 inhibited TEAD4 palmitoylation through direct interaction. Acyl-protein thioesterase 2 (APT2) is known as a major depalmitoylase of TEAD family proteins (Kim and Gumbiner, 2019). We found that the protein level of APT2 was not affected by the expression of NF2 (**Figure supplement 4A**), which excluded the possibility that NF2 reduced TEAD4 palmitoylation through APT2.

The depalmitoylation has been shown to trigger the degradation of TEAD protein mediated by E3 ligase CHIP (Kim and Gumbiner, 2019). The *in vitro* ubiquitination assays confirmed that non-palmitoylated mutant TEAD4-2CS (C335S/C367S) exhibited much higher ubiquitination levels than TEAD4-WT (**Figure 5F**). As positively relevant, TEAD4-2CS also exhibited stronger binding to NF2 than TEAD4-WT (**Figure 5G**), indicating that NF2 preferentially bound to non-palmitoylated form of TEAD4 and triggered its ubiquitination. Thus, NF2 inhibits TEAD4 palmitoylation and presumably causes the sequential ubiquitination of TEAD4.

### TEAD4 interaction is required for NF2 function to suppress cell proliferation

We then explored whether the direct interaction between NF2 and TEAD4 contributes to the tumor suppressor function of NF2. The BrdU incorporation assay was performed in *NF2* KO HEK293A cells to examine cell proliferation rates. In comparison with control cells, BrdU incorporation efficiency was dramatically increased in *NF2* KO cells, and significantly decreased with NF2-WT expression to a similar level with control cells (**Figure 6A and 6B**), confirming the tumor suppression role of NF2. However, the cells expressing NF2-5A mutant kept a higher incorporation efficiency of BrdU around 90% of *NF2* KO cells (**Figure 6A and 6B**). NF2 suppressed cell proliferation as the tumor suppressor did, whereas the TEAD4-binding deficient mutant of NF2 lacks the suppressor function, indicating that TEAD4 interaction is required for NF2 function to suppress cell proliferation.

**Figure 6.**
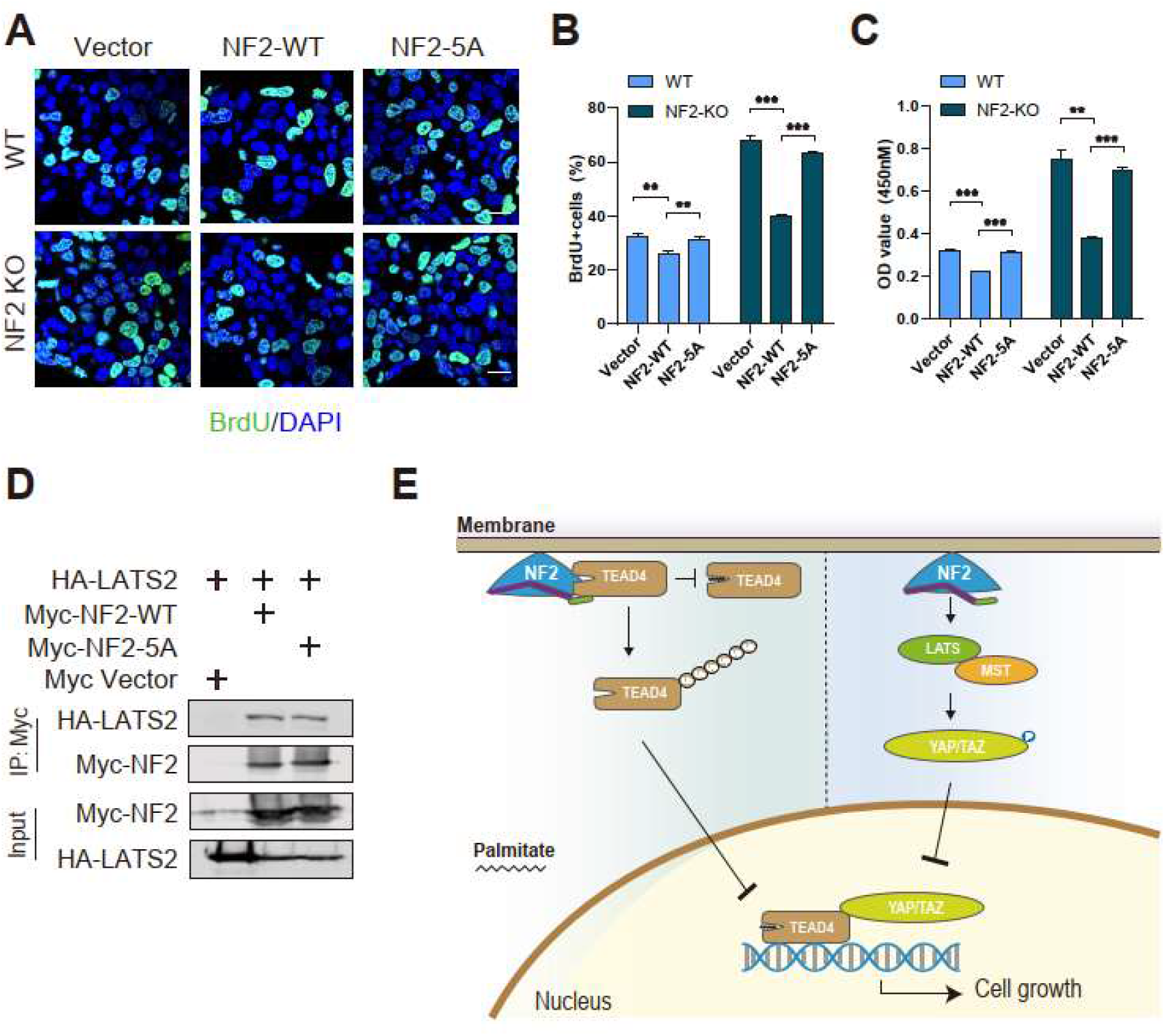
TEAD4interaction is required for NF2 function to suppress tumor cell proliferation. (A) WT and *NF2* KO HEK293A cells transfected with vector or the NF2-WT/5A were subjected to BrdU incorporation assay. Scale bar = 20 μm. (B) The percentage of BrdU-positive cells from (A) was quantified. Mean ± s.e.m, N = 3, \*\**P*< 0.01, \*\*\**P*< 0.001. (C) WT and *NF2* KO HEK293A cells transfected with vector or the NF2-WT/5A were subjected to cell counting kit-8 (CCK-8), and OD 450nm were quantified. Mean ± s.e.m.\*\**P*< 0.01, \*\*\**P*< 0.001. (D) Co-immunoprecipitation experiment of Myc-tagged NF2 WT and mutants with HA-tagged LATS2 was performed in *NF2* KO HEK293A cells. Cell lysates were treated to anti-Myc beads and immunoblotted with indicated antibodies. (E) The working model for the molecular mechanism of NF2 function to suppress tumor cell proliferation through LATS1/2 (right) and directly through TEAD4 (left). The straightforward mechanism, which NF2 directly binds to and down-regulates TEAD4 activity, could complement the classical regulation through Hippo pathway. **Figure 6-source data 1**. Source data for quantifications graphed in ***Figure 6B***. **Figure 6-source data 2**. Source data for quantifications graphed in ***Figure 6C***. **Figure 6-source data 3**. Whole uncropped blots represented in ***Figure 6D***. HA-Lats and Myc-NF2 protein levels with Co-immunoprecipitation assay in *NF2* KO HEK293A cells.

The cell proliferation suppression by NF2 WT and mutant in *NF2* KO HEK293A cells was further measured by CCK-8 cell viability assay. Similar to the results from BrdU incorporation experiment, NF2-WT robustly decreased cell viability, while NF2-5A dramatically restored cell viability (**Figure 6C**). Since NF2 can interact with and activate LATS1/2 in the Hippo signaling, we then examined whether NF2-5A mutant disrupt the interaction with LATS. Co-immunoprecipitation assay in *NF2* KO cells showed that NF2-5A mutant exhibited similar binding activity to LATS2, compared with NF2-WT (**Figure 6D**), indicating that the suppression defect of cell proliferation induced by NF2-5A is not related to LATS1/2 activation. Taken together, TEAD4 binding is required for NF2 function to suppress cell proliferation, and is presumably caused by inhibiting TEAD4 palmitoylation.

## Discussion

As a tumor suppressor, NF2 senses cell-cell contact and regulates the Hippo pathway by activating LATS1/2 kinases, resulting in phosphorylation and cytoplasmic retention of YAP. Phosphorylated -YAP could not form complex with transcription factor TEAD in the nucleus, thereby inhibiting cell proliferation and suppressing tumor growth (Meng et al., 2016; Morrison et al., 2001; Okada et al., 2005; Yin et al., 2013). In contrast to the classic model of NF2, our finding proposed a straightforward regulation mechanism that NF2 directly associates with TEAD4 to promote the cytoplasmic retention and inhibit palmitoylation of TEAD4, resulting in dysfunction of TEAD4 and cell proliferation suppression (**Figure 6E**). We further validated 5 key residues of NF2 is required for TEAD4 interaction and cell proliferation suppression. The missense mutations in these sites, such as L297V, H304Y, and F591L, are also found in various cancers (Bonilla et al., 2016; Zehir et al., 2017). NF-5A mutant lost the binding ability to TEAD4, but still bound to LATS2, which indicates that the function deficient of NF2-5A is because of the binding defect to TEAD instead of the classical Hippo pathway. This straightforward regulation would shed light on additional mechanism of how tumor suppressor NF2 functions, and also complement the regulation from LATS1/2 of Hippo pathway.

As transcription factor, TEAD family proteins form transcriptional complex with the major co-activators YAP/TAZ to activate the transcription of important target genes and promote cell proliferation and organ growth (Yu et al., 2015). Besides that, The inhibitory binding partners of TEADs, such as VGLL4, compete with YAP/TAZ and inhibit transcription activity of TEAD4 to suppress cell proliferation and tumor growth in multiple cancers (Jiao et al., 2014; Zhang et al., 2014). Our finding that tumor suppressor NF2 inhibits TEAD4 palmitoylation via direct interaction, proposed a new role for the classical protein NF2 as the inhibitory binding partner of TEAD4, which might update the functional cognition of NF2 in the Hippo pathway.

Palmitoylation is essential for protein stability and transcription activity of TEADs (Chan et al., 2016; Noland et al., 2016), and NF2 has been shown to decrease the mRNA levels of fatty acid synthase (FASN) and induce depalmitoylation of TEADs (Kim and Gumbiner, 2019). Alternatively, we found that palmitoylation of TEAD4 could also be inhibited by NF2 via direct interaction, which might reflect a novel role of NF2 in regulating palmitoylation and homeostasis of TEAD protein in cells.

The nuclear localization is also required for TEADs transcription activity. Early studies have reported the cytoplasmic translocation of TEADs induced by cell density and p38 (Lin et al., 2017). As a membrane-associated protein, NF2 has been shown to recruit and activate LATS1/2 kinases at plasma membrane (Yin et al., 2013). Here, we showed that NF2 also induced cytoplasmic translocation of TEAD4 via direct protein-protein interactions. The translocations of both LATS1/2 and TEADs induced by NF2 reach the same goal preventing TEADs transcription activity, highly suggested that this straightforward regulation could complement the function of Hippo pathway.

In summary, we identified the direct link and physical interaction between NF2 and TEAD4, which are important for NF2 function as tumor suppressor and TEADs protein homeostasis.

## Acknowledgements

We thank Faxing Yu (Children’s Hospital of Fudan University) for the *NF2* KO HEK293A cell line. This work was supported by National Key R&D Program of China [2019YFA0802000 to L.Z.], the National Natural Science Foundation of China [31970672 to Xiaojing He, 32000496 to Liqiao Hu and 32030025 and 31625017 to L. Z.], Shanghai Leading Talents Program to L.Z. and China Postdoctoral Science Foundation [2019M662588 to Liqiao Hu].

## Author Contributions

Conceptualization: LQ.H., L.Z. and XJ.H; Methodology: LQ.H., MY.W., LL.H., L.Y., B.Z., L.Z. and XJ.H.; Investigation: LQ.H., MY.W., LL.H. and L.Y.; Writing-Original Draft: LQ.H. and MY.W.; Writing– Review & Editing: LL.H., B.Z., L.Z. and XJ.H.; Funding Acquisition: LQ.H., L.Z. and XJ.H.; Resources: L.Z. and XJ.H.; Supervision: XJ.H.

## Materials and methods

### Protein purification

Human TEAD4 YBD domain (217-434) was cloned into a PET-28a vector with an N-terminal SUMO and 6 ×His tag and a PGEX-6P-1 vector with an N-terminal GST tag, respectively. SUMO-tagged and GST-tagged TEAD4-YBD were expressed in *Escherichia coli* BL21 (DE3) cells and purified using the Ni^2+^-NTA agarose resin (GE Healthcare) or GST agarose resin (GE Healthcare), and purified via size-exclusion chromatography using a Superdex 200 column (GE Healthcare). Purified SUMO-TEAD4 and GST-TEAD4 were concentrated to 2 mg/mL in a buffer containing 20 mM Tris (pH 8.0), 150 mM NaCl, and 1 mM DTT.

For TEAD4-YBD alone, GST-TEAD4-YBD was purified using GST agarose resin (GE Healthcare). GST tag was removed with 3C protease overnight at 4 °C. The resin was collected using the tag-free with 3C protease overnight at 4°C. The eluted TEAD4-YBDprotein was purified via size-exclusion chromatography using a Superdex 200 column (GE Healthcare). Purified TEAD4-YBD was concentrated to 2 mg/mL in a buffer containing 20 mM Tris (pH 8.0), 150 mM NaCl, and 1 mM DTT.

NF2(18-595) was cloned into a PMAL-3C vector with an N-terminal MBP tag. MBP-NF2 was purified using amylose resin (New England Biolabs) and eluted using 10mM maltose (BioFroxx). MBP-NF2 was purified via size exclusion chromatography using a Superdex 200 column (GE Healthcare) in a buffer containing 20 mM Tris pH 8.0, 150 mM NaCl, and 1 mM DTT. Human CHIP and Hsp70 were cloned into a PET-28a vector with an N-terminal SUMO tag. SUMO-CHIP and SUMO-Hsp70 were purified using the Ni^2+^-NTA agarose resin (GE Healthcare) and eluted with 200mM imidazole (Sigma-Aldrich). The proteins were then purified with a Superdex 200 column (GE Healthcare) in a buffer containing 20 mM Tris (pH 8.0), 150 mM NaCl, and 1 mM DTT.

### Cell culture and transfection

HEK293T, HEK293A, HeLa, and MCF-10A cells were cultured in DMEM medium (Gibco) supplemented with 10% fetal bovine serum (FBS, Gibco) and 1% penicillin/streptomycin. NCI-H226 cells were cultured in the Roswell Park Memorial Institute-1640 medium (Gibco) supplemented with 10% FBS (Gibco) and 1% penicillin/streptomycin. Plasmids were transfected using the HighGene Transfection Reagent (ABclonal). HeLa cells were transfected using Lipofectamine 2000 (Invitrogen). The sequences of siRNAs used in this study are as follows:

*siNF2-1:* CCGUGAGGAUCGUCACCAUTT

*siNF2-2:* GGUACUGGAUCAUGAUGUUTT

*siNF2-3:* GGAAUGAAAUCCGAAACAUTT

### Protein immunoprecipitation

HEK293T cells were transiently transfected with Myc-NF2 and Flag-TEAD4. The cells were lysed in a lysis buffer (20 mM Tris 7.5, 150 mM NaCl, 1 mM EDTA, 1% Triton, and phosphatase inhibitor cocktail) for 30 min at 0°C. The supernatant was incubated with red anti-FLAG beads (Millipore) or Protein-A Magnetic beads (Bio-Rad) and Myc antibody (Cell Signaling Technology) overnight at 4°C. The proteins on the beads were subjected to SDS-PAGE and analyzed via western blotting.

### *In vitro* protein-binding assay

Recombinant GST-NF2 was bound to a GST resin (GE Healthcare), and MBP-NF2 was bound to an MBP resin (New England Biolabs) in PBS for 1 h at 4°C. After washing, the resin was incubated with purified SUMO-TEAD4 in PBS for 1h at 4°C and washed four times. Proteins retained on the beads were analyzed using SDS-PAGE and western blotting. SUMO-TEAD4 was detected using an antibody against 6 × His.

### *In vitro* palmitoylation assay

Recombinant TEAD4 protein (500 ng) was incubated with 1 mM alkyne palmitoyl-CoA (Cayman Chemical) for 0.5 h in 20mM Tris 8.0 and 100mM NaCl. Click reaction with biotin-azide (Sigma-Aldrich) was performed for 1h at 25°C. The reactions were stopped using 2XSDS sample buffer, followed by SDS-PAGE analysis. Biotinylated TEAD4 was detected using streptavidin-IRDye (LI-COR).

### *In vivo* palmitoylation assay

Myc-NF2 was transfected into the cells and 48 h after transfection, the cells were collected and subjected to the CAPTUREome S-Palmitoylated Protein Kit (Badrilla). Briefly, the cells were lysed and blocked with the blocking buffer at 40°C for 4 h. The mixture was then subjected to ice-cold acetone precipitation. The precipitate was re-dissolved in the binding buffer and incubated with the thioester cleavage reagent and capture resin for 2 h. After washing, the capture resin was subjected to SDS-PAGE and analyzed via western blotting.

### *In vitro* ubiquitination assay

Each in vitro ubiquitination reaction was performed using 0.5uM E1, 4uM UbcH5b, 2uM CHIP, 1uM Hsp70, 10uM ubiquitin, and 1uM recombinant GST-WT/2CS TEAD4 for 60 min at 37°C in 20mM Tris8.0, 100mM NaCl, 5mM ATP, 2.5mM MgCl_2_, and 1mM DTT. Ubiquitination reactions were stopped using 2XSDS sample buffer, followed by detection via western blotting with the GST antibody (ABclonal).

### Real-time PCR

Total RNA was extracted using the TRIzol reagent (Invitrogen), and reverse transcription (RT) was performed using the iScript Reverse Transcription Supermix (Bio-Rad). Real-time RT-PCR analysis was performed using SYBR Green Realtime PCR Master Mix (Toyobo) with the Applied Biosystems Step Two Real-Time PCR System (Applied Biosystems). GAPDH was used as a control. The standard comparative CT quantization method was used to analyze the RT-PCR results.

Primers for RT-PCR are as following:

*TEAD1* F: ATGGAAAGGATGAGTGACTCTGC

*TEAD1* R: TCCCACATGGTGGATAGATAGC

*TEAD2* F: CTTCGTGGAACCGCCAGAT

*TEAD2* R: GGAGGCCACCCTTTTTCTCA

*TEAD3* F: TCATCCTGTCAGZCGAGGG

*TEAD3* R: TCTTCCGAGCTAGAACCTGTATG

*TEAD4* F: GAACGGGGACCCTCCAATG

*TEAD4* R: GCGAGCATACTCTGTCTCAAC

*YAP* F: CACAGCATGTTCGAGCTCAT

*YAP* R: GATGCTGAGCTGTGGGTGTA

*NF2* F: TGCGAGATGAAGTGGAAAGG

*NF2* R: GCCAAGAAGTGAAAGGTGAC

### BiFC assay

Full-length YFP (1-238) was divided into two insertions: nYFP (1-154) and cYFP (155-238) in this study. pcDNA3.1-NF2/TEAD4/LATS vectors were receptions for C-terminal n/c-YFP fragments with HindIII and BamHIsites.HEK-293T cells plated in a 6-well plate for 24 h, and transfected with 800 ng nYFP- and 800 ng cYFP-tagged NF2/TEAD4?LATS constructs. The cells were treated at low temperature (30 °C) for 6 h for fluorophore maturation, and after 48 h, fluorescence was determined via flow cytometry using BD FACS Calibur (BD Biosciences) or observed under a confocal laser scanning microscope (Olympus).

### Immunofluorescent microscopy

NF2 KO HEK293A cells on coverslips were transfected with Myc-NF2 WT or mutant for 48h at 37°C.Cells were fixed in 4% paraformaldehyde (aladin) for 30 min followed by permeabilization with 0.1% TritonX-100 (aladin) for 30 mins. Cells were blocked in 3% BSA for 1 h and incubated overnight at 4°C in primary antibodies diluted in 3% BSA. Secondary antibodies were diluted in 3% BSA and incubated for 1 h. Then cells were stained with DAPI (Beyotime).

### BrdU Incorporation assay

WT and *NF2* KOHEK293A cells on coverslips were transfected with Myc-NF2 WT or mutant and after 36 h incubated with 10 uM BrdU (Beyotime) for 6 h at 37°C. Cells were fixed with 4% paraformaldehyde for 30 min and washed with PBS with 1% Triton X-100 for 30 min. Then cells were incubated with 2N HCl for 30 min at room temperature. After washing with PBS, cells were blocked with PBS containing 1% Triton X-100 and 3% BSA. Cells were incubated with the primary antibodies against BrdU (ABCam) overnight at 4°C. and incubated with the Alexa Fluor 488 dye-conjugated secondary antibodies (Invitrogen) for 1 h in the dark and then stained with DAPI (Beyotime).

### Cell counting kit-8 (CCK-8) assay

Cells were seeded into 96-well plates and transfected with Myc-NF2 WT or mutant. After 48 h, cell counting kit-8 (Biosharp) was used to detect cell viability. The cells in each well were incubated with 10 ul CCK-8 solution at 37°C for 1 h. The absorbance at 450 nm was detected using a plate reader.

## Supplemental material

**Figure supplement 1.**
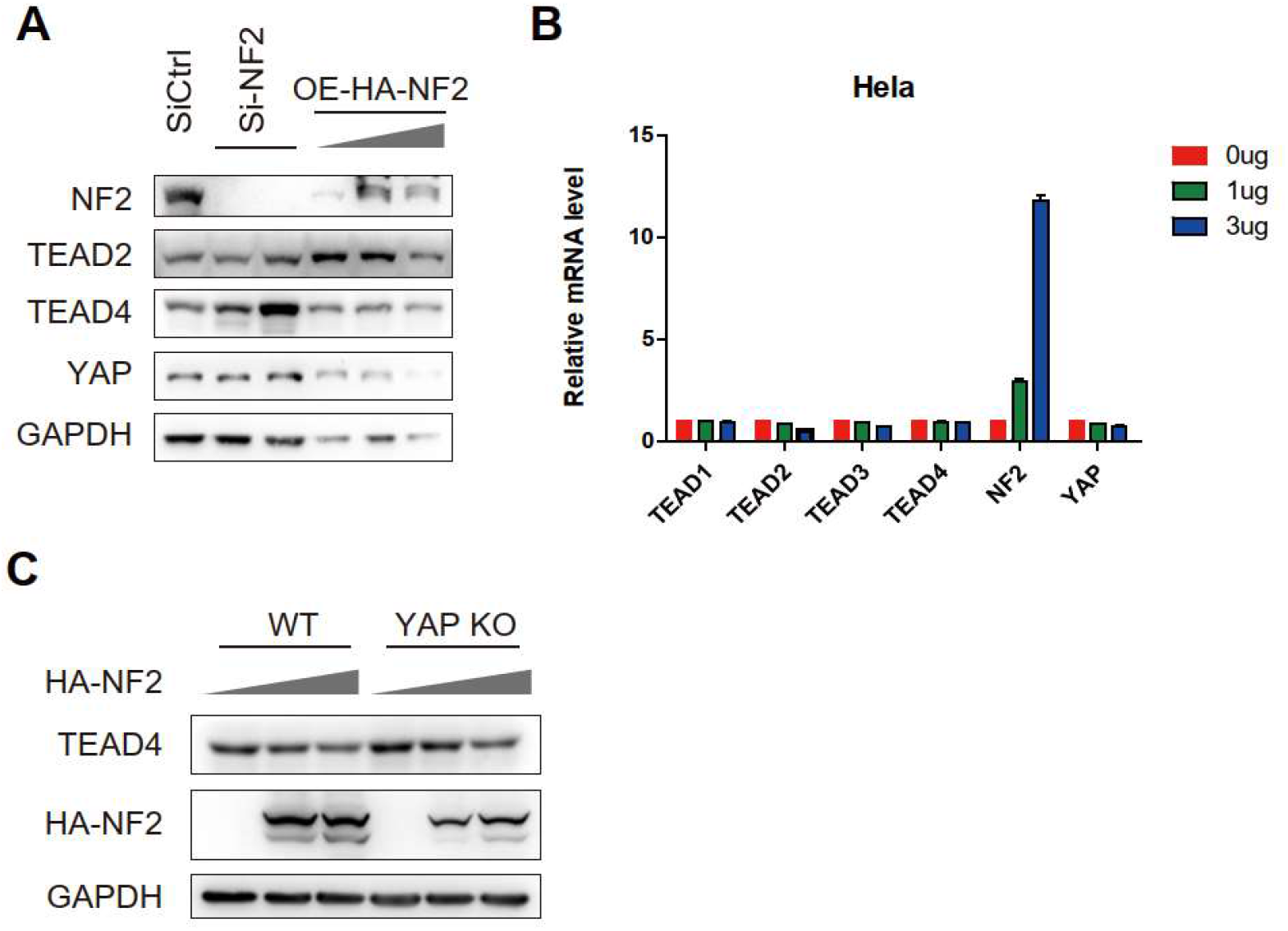
NF2 decreases the protein levels of TEADs. (A) Protein levels of TEAD2/4 and YAP were determined by western blotting in MCF-10A cells with overexpression of HA-NF2 or siNF2 treatment. (B) RT-PCR analysis of TEAD1/2/3/4 and YAP mRNA levels were performed in Hela cells with overexpression of NF2. (C) Protein level of TEAD4 was determined by western blotting in WT or YAP-KO HEK293T cells with overexpression of HA-NF2. **Figure supplement 1-source data 1**. Whole uncropped blots represented in ***Figure supplement 1A***. NF2, TEAD2, TEAD4, YAP and GADPH protein levels in MCF-10A cells. **Figure supplement 1-source data 2**. Whole uncropped blots represented in ***Figure supplement 1C***. NF2, TEAD4 and GADPH protein levels in WT or YAP-KO HEK293T cells.

**Figure supplement 2.**
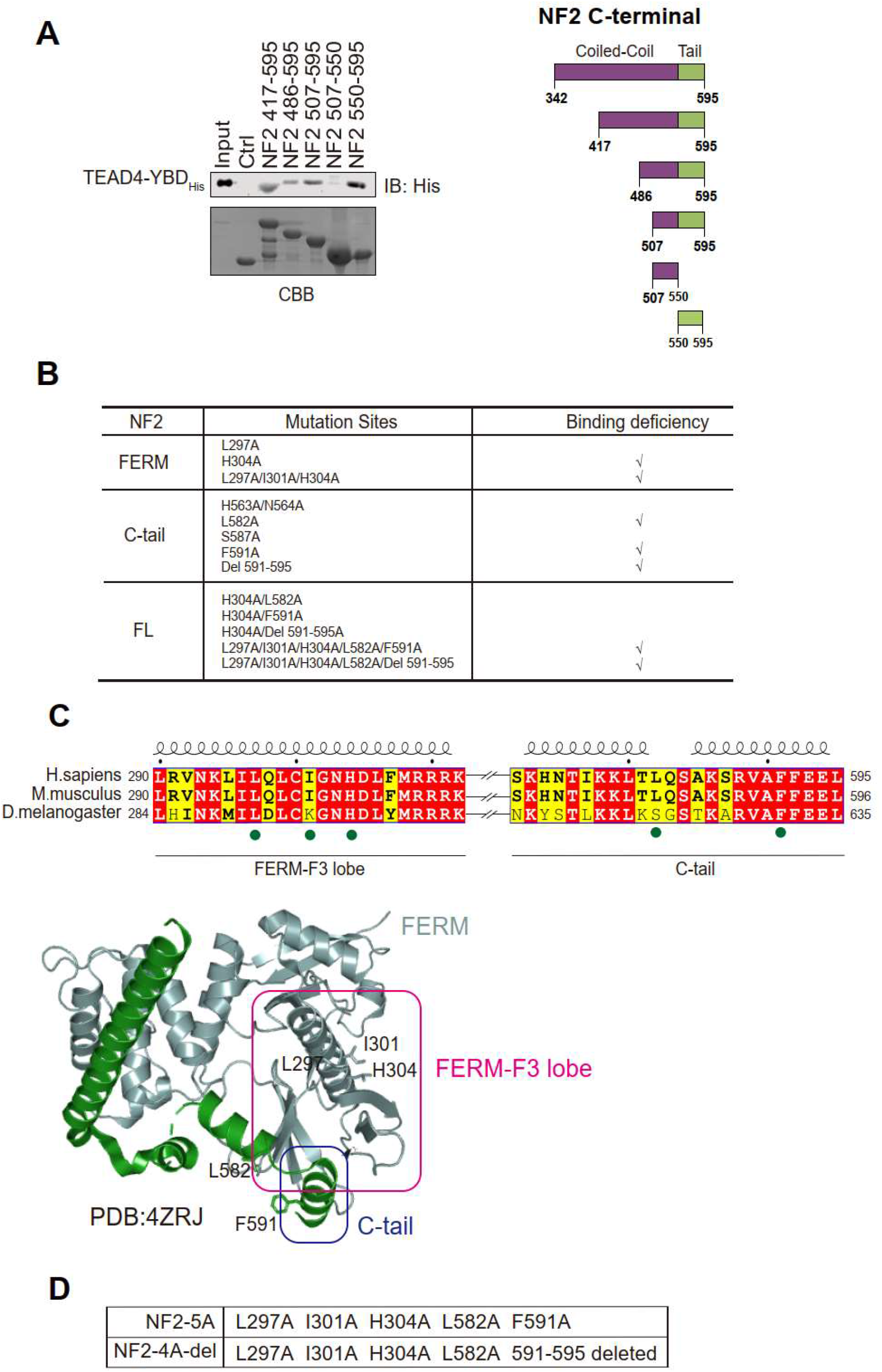
Characterization of the interaction between NF2 and TEAD4. (A) The GST-pull down assay was performed to assess the interaction between TEAD4-YBD and NF2 C-terminal truncations. CBB, Coomassie brilliant blue. The schematic views of NF2 C-terminal truncations showed in right panel. Related to Figure 2. (B) The indicated residues on the surface of NF2 were screened for the interaction with TEAD4 by GST binding assay. N/A, not available. Related to Figure 3. (C) The five key residues of NF2 were pinpointed to mediate interaction with TEAD4. (D)Table of NF2 mutation sites applied in this study. **Figure supplement 2-source data 1**. Whole SDS-PAGE images and uncropped blots represented in ***Figure supplement 2A***. TEAD4-YBD_His_ and GST-NF2 fragment protein levels in GST pull-down assay.

**Figure supplement 3.**
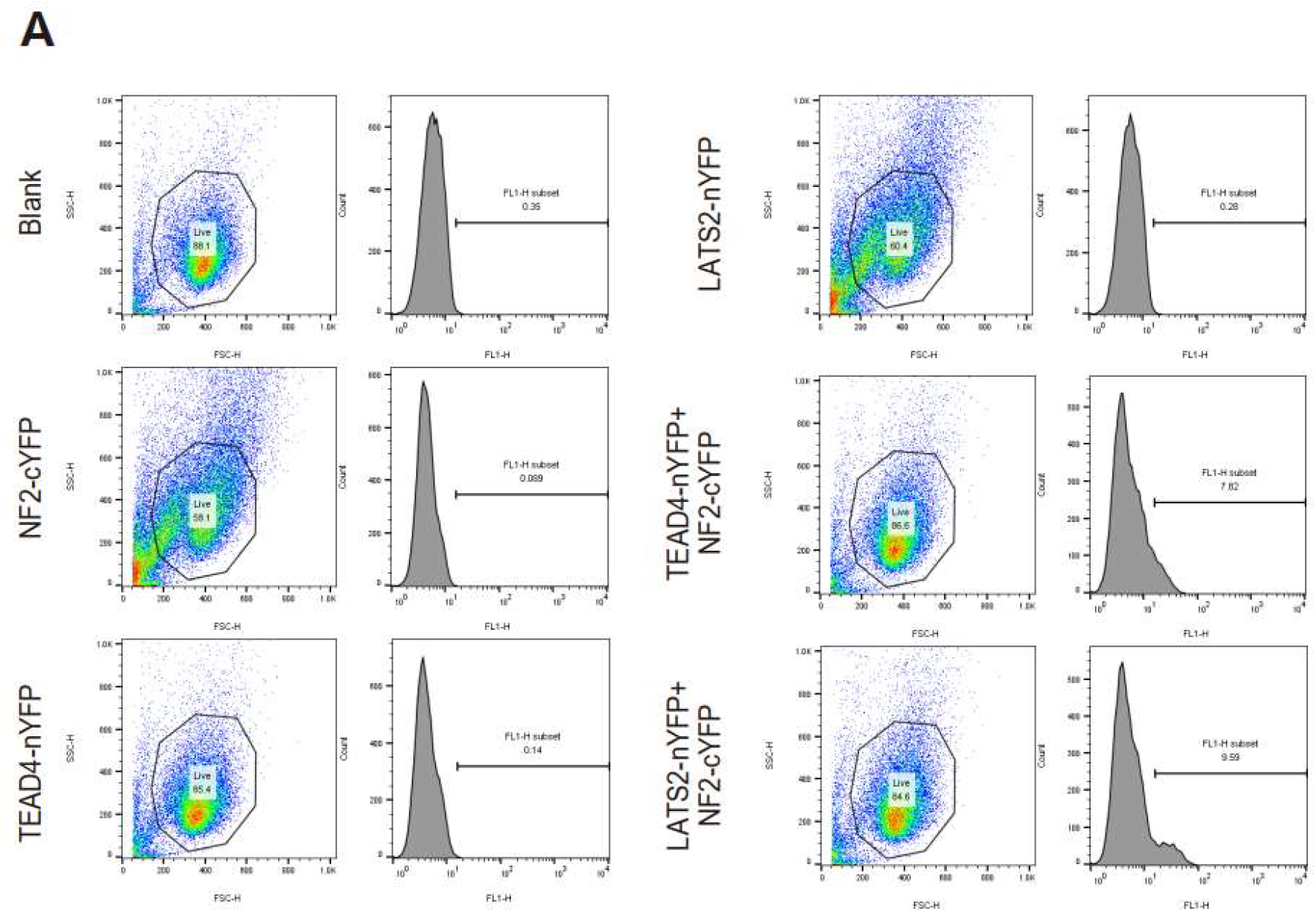
The flow cytometry to measure fluorescence signals of indicated cells from BiFC assay. (A) Fluorescence signals of the cells used in BiFC assay were sequentially measured and quantified by flow cytometry. Gating strategies used for flow cytometry and cells were selected in the FSC-H/SSC-H dot plot to remove debris. Quantification of signals from flow cytometry showed in Figure 4B.

**Figure supplement 4.**
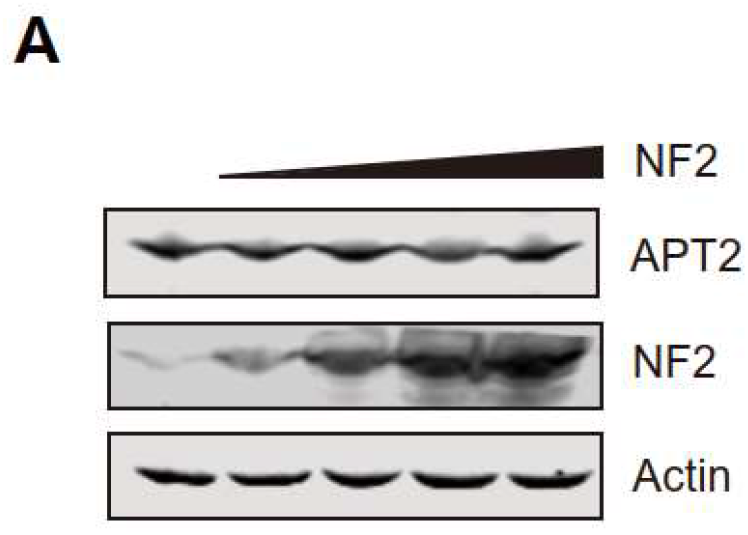
The overexpression of NF2 in cells does not affect APT2 protein levels. (A) NCI-H226 cells transfected Myc-NF2 to determine the protein levels of APT2 via immunoblotting. **Figure supplement 4-source data 1**. Whole uncropped blots represented in ***Figure supplement 4A***. APT2, NF2 and beta-actin protein levels in NCI-H226 cells.

## References

Bonilla X, Parmentier L, King B, Bezrukov F, Kaya G, Zoete V, Seplyarskiy VB, Sharpe HJ, McKee T, Letourneau A, Ribaux PG, Popadin K, Basset-Seguin N, Ben Chaabene R, Santoni FA, Andrianova MA, Guipponi M, Garieri M, Verdan C, Grosdemange K, Sumara O, Eilers M, Aifantis I, Michielin O, de Sauvage FJ, Antonarakis SE, Nikolaev SI. 2016. Genomic analysis identifies new drivers and progression pathways in skin basal cell carcinoma. Nat Genet 48:398–406. doi:10.1038/ng.3525

Bum-Erdene K, Zhou D, Gonzalez-Gutierrez G, Ghozayel MK, Si Y, Xu D, Shannon HE, Bailey BJ, Corson TW, Pollok KE, Wells CD, Meroueh SO. 2019. Small-Molecule Covalent Modification of Conserved Cysteine Leads to Allosteric Inhibition of the TEAD·Yap Protein-Protein Interaction. Cell Chemical Biology 26:378-389.e13. doi:10.1016/j.chembiol.2018.11.010

Chan P, Han X, Zheng B, DeRan M, Yu J, Jarugumilli GK, Deng H, Pan D, Luo X, Wu X. 2016. Autopalmitoylation of TEAD proteins regulates transcriptional output of the Hippo pathway. Nat Chem Biol 12:282–289. doi:10.1038/nchembio.2036

Chen H, Xue L, Wang H, Wang Z, Wu H. 2017. Differential NF2 Gene Status in Sporadic Vestibular Schwannomas and its Prognostic Impact on Tumour Growth Patterns. Sci Rep 7:5470. doi:10.1038/s41598-017-05769-0

Cheng JQ, Lee W-C, Klein MA, Cheng GZ, Jhanwar SC, Testa JR. 1999. Frequent mutations of NF2 and allelic loss from chromosome band 22q12 in malignant mesothelioma: Evidence for a two-hit mechanism ofNF2 inactivation. Genes Chromosom Cancer 24:238–242. doi:10.1002/(SICI)1098-2264(199903)24:3<238::AID-GCC9>3.0.CO;2-M

Chinthalapudi K, Mandati V, Zheng J, Sharff AJ, Bricogne G, Griffin PR, Kissil J, Izard T. 2018. Lipid binding promotes the open conformation and tumor-suppressive activity of neurofibromin 2. Nat Commun 9:1338. doi:10.1038/s41467-018-03648-4

Cooper J, Giancotti FG. 2014. Molecular insights into NF2 /Merlin tumor suppressor function. FEBS Letters 588:2743–2752. doi:10.1016/j.febslet.2014.04.001

Deng X, Fang L. 2018. VGLL4 is a transcriptional cofactor acting as a novel tumor suppressor via interacting with TEADs. Am J Cancer Res 8:932–943.

Dey A, Varelas X, Guan K-L. 2020. Targeting the Hippo pathway in cancer, fibrosis, wound healing and regenerative medicine. Nat Rev Drug Discov 19:480–494. doi:10.1038/s41573-020-0070-z

Forrester MT, Hess DT, Thompson JW, Hultman R, Moseley MA, Stamler JS, Casey PJ. 2011. Site-specific analysis of protein S-acylation by resin-assisted capture. J Lipid Res 52:393–398. doi:10.1194/jlr.D011106

Gupta MP, Kogut P, Gupta M. 2000. Protein kinase-A dependent phosphorylation of transcription enhancer factor-1 represses its DNA-binding activity but enhances its gene activation ability. Nucleic Acids Res 28:3168–3177. doi:10.1093/nar/28.16.3168

Harvey KF, Pfleger CM, Hariharan IK. 2003. The Drosophila Mst ortholog, hippo, restricts growth and cell proliferation and promotes apoptosis. Cell 114:457–467. doi:10.1016/s0092-8674(03)00557-9

He L, Yuan L, Sun Y, Wang P, Zhang H, Feng X, Wang Z, Zhang W, Yang C, Zeng YA, Zhao Y, Chen C, Zhang L. 2019. Glucocorticoid Receptor Signaling Activates TEAD4 to Promote Breast Cancer Progression. Cancer Res 79:4399–4411. doi:10.1158/0008-5472.CAN-19-0012

Hong AW, Meng Z, Plouffe SW, Lin Z, Zhang M, Guan K-L. 2020. Critical roles of phosphoinositides and NF2 in Hippo pathway regulation. Genes Dev 34:511–525. doi:10.1101/gad.333435.119

Huang J, Wu S, Barrera J, Matthews K, Pan D. 2005. The Hippo Signaling Pathway Coordinately Regulates Cell Proliferation and Apoptosis by Inactivating Yorkie, the Drosophila Homolog of YAP. Cell 122:421–434. doi:10.1016/j.cell.2005.06.007

Huh H, Kim D, Jeong H-S, Park H. 2019. Regulation of TEAD Transcription Factors in Cancer Biology. Cells 8:600. doi:10.3390/cells8060600

Jiang SW, Dong M, Trujillo MA, Miller LJ, Eberhardt NL. 2001. DNA binding of TEA/ATTS domain factors is regulated by protein kinase C phosphorylation in human choriocarcinoma cells. J Biol Chem 276:23464–23470. doi:10.1074/jbc.M010934200

Jiao S, Li C, Hao Q, Miao H, Zhang L, Li L, Zhou Z. 2017. VGLL4 targets a TCF4–TEAD4 complex to coregulate Wnt and Hippo signalling in colorectal cancer. Nat Commun 8:14058. doi:10.1038/ncomms14058

Jiao S, Wang H, Shi Z, Dong A, Zhang W, Song X, He F, Wang Y, Zhang Z, Wang W, Wang X, Guo T, Li P, Zhao Y, Ji H, Zhang L, Zhou Z. 2014. A Peptide Mimicking VGLL4 Function Acts as a YAP Antagonist Therapy against Gastric Cancer. Cancer Cell 25:166–180. doi:10.1016/j.ccr.2014.01.010

Kalamarides M. 2002. Nf2 gene inactivation in arachnoidal cells is rate-limiting for meningioma development in the mouse. Genes & Development 16:1060–1065. doi:10.1101/gad.226302

Kim N-G, Gumbiner BM. 2019. Cell contact and Nf2/Merlin-dependent regulation of TEAD palmitoylation and activity. Proc Natl Acad Sci USA 116:9877–9882. doi:10.1073/pnas.1819400116

Li W, You L, Cooper J, Schiavon G, Pepe-Caprio A, Zhou L, Ishii R, Giovannini M, Hanemann CO, Long SB, Erdjument-Bromage H, Zhou P, Tempst P, Giancotti FG. 2010. Merlin/NF2 Suppresses Tumorigenesis by Inhibiting the E3 Ubiquitin Ligase CRL4DCAF1 in the Nucleus. Cell 140:477–490. doi:10.1016/j.cell.2010.01.029

Li Y, Zhou H, Li F, Chan SW, Lin Z, Wei Z, Yang Z, Guo F, Lim CJ, Xing W, Shen Y, Hong W, Long J, Zhang M. 2015. Angiomotin binding-induced activation of Merlin/NF2 in the Hippo pathway. Cell Res 25:801–817. doi:10.1038/cr.2015.69

Lin KC, Moroishi T, Meng Z, Jeong H-S, Plouffe SW, Sekido Y, Han J, Park HW, Guan K-L. 2017. Regulation of Hippo pathway transcription factor TEAD by p38 MAPK-induced cytoplasmic translocation. Nat Cell Biol 19:996–1002. doi:10.1038/ncb3581

Liu X, Li H, Rajurkar M, Li Q, Cotton JL, Ou J, Zhu LJ, Goel HL, Mercurio AM, Park J-S, Davis RJ, Mao J. 2016. Tead and AP1 Coordinate Transcription and Motility. Cell Reports 14:1169–1180. doi:10.1016/j.celrep.2015.12.104

Meng Z, Moroishi T, Guan K-L. 2016. Mechanisms of Hippo pathway regulation. Genes Dev 30:1–17. doi:10.1101/gad.274027.115

Mesrouze Y, Meyerhofer M, Bokhovchuk F, Fontana P, Zimmermann C, Martin T, Delaunay C, Izaac A, Kallen J, Schmelzle T, Erdmann D, Chène P. 2017. Effect of the acylation of TEAD4 on its interaction with co-activators YAP and TAZ: TEAD Acylation. Protein Science 26:2399–2409. doi:10.1002/pro.3312

Morrison H, Sherman LS, Legg J, Banine F, Isacke C, Haipek CA, Gutmann DH, Ponta H, Herrlich P. 2001. The NF2 tumor suppressor gene product, merlin, mediates contact inhibition of growth through interactions with CD44. Genes Dev 15:968–980. doi:10.1101/gad.189601

Noland CL, Gierke S, Schnier PD, Murray J, Sandoval WN, Sagolla M, Dey A, Hannoush RN, Fairbrother WJ, Cunningham CN. 2016. Palmitoylation of TEAD Transcription Factors Is Required for Their Stability and Function in Hippo Pathway Signaling. Structure 24:179–186. doi:10.1016/j.str.2015.11.005

Okada T, Lopez-Lago M, Giancotti FG. 2005. Merlin/NF-2 mediates contact inhibition of growth by suppressing recruitment of Rac to the plasma membrane. J Cell Biol 171:361–371. doi:10.1083/jcb.200503165

Ota M, Sasaki H. 2008. Mammalian Tead proteins regulate cell proliferation and contact inhibition as transcriptional mediators of Hippo signaling. Development 135:4059–4069. doi:10.1242/dev.027151

Pan D. 2010. The hippo signaling pathway in development and cancer. Dev Cell 19:491–505. doi:10.1016/j.devcel.2010.09.011

Pobbati AV, Han X, Hung AW, Weiguang S, Huda N, Chen G-Y, Kang C, Chia CSB, Luo X, Hong W, Poulsen A. 2015. Targeting the Central Pocket in Human Transcription Factor TEAD as a Potential Cancer Therapeutic Strategy. Structure 23:2076–2086. doi:10.1016/j.str.2015.09.009

Sher I, Hanemann CO, Karplus PA, Bretscher A. 2012. The tumor suppressor merlin controls growth in its open state and is converted by phosphorylation to a less-active more-closed state. Developmental cell 22:703. doi:10.1016/j.devcel.2012.03.008

Wu S, Huang J, Dong J, Pan D. 2003. hippo Encodes a Ste-20 Family Protein Kinase that Restricts Cell Proliferation and Promotes Apoptosis in Conjunction with salvador and warts. Cell 114:445–456. doi:10.1016/S0092-8674(03)00549-X

Yin F, Yu J, Zheng Y, Chen Q, Zhang N, Pan D. 2013. Spatial Organization of Hippo Signaling at the Plasma Membrane Mediated by the Tumor Suppressor Merlin/NF2. Cell 154:1342–1355. doi:10.1016/j.cell.2013.08.025

Yu F-X, Zhao B, Guan K-L. 2015. Hippo Pathway in Organ Size Control, Tissue Homeostasis, and Cancer. Cell 163:811–828. doi:10.1016/j.cell.2015.10.044

Zehir A, Benayed R, Shah RH, Syed A, Middha S, Kim HR, Srinivasan P, Gao J, Chakravarty D, Devlin SM, Hellmann MD, Barron DA, Schram AM, Hameed M, Dogan S, Ross DS, Hechtman JF, DeLair DF, Yao J, Mandelker DL, Cheng DT, Chandramohan R, Mohanty AS, Ptashkin RN, Jayakumaran G, Prasad M, Syed MH, Rema AB, Liu ZY, Nafa K, Borsu L, Sadowska J, Casanova J, Bacares R, Kiecka IJ, Razumova A, Son JB, Stewart L, Baldi T, Mullaney KA, Al-Ahmadie H, Vakiani E, Abeshouse AA, Penson AV, Jonsson P, Camacho N, Chang MT, Won HH, Gross BE, Kundra R, Heins ZJ, Chen H-W, Phillips S, Zhang H, Wang J, Ochoa A, Wills J, Eubank M, Thomas SB, Gardos SM, Reales DN, Galle J, Durany R, Cambria R, Abida W, Cercek A, Feldman DR, Gounder MM, Hakimi AA, Harding JJ, Iyer G, Janjigian YY, Jordan EJ, Kelly CM, Lowery MA, Morris LGT, Omuro AM, Raj N, Razavi P, Shoushtari AN, Shukla N, Soumerai TE, Varghese AM, Yaeger R, Coleman J, Bochner B, Riely GJ, Saltz LB, Scher HI, Sabbatini PJ, Robson ME, Klimstra DS, Taylor BS, Baselga J, Schultz N, Hyman DM, Arcila ME, Solit DB, Ladanyi M, Berger MF. 2017. Mutational landscape of metastatic cancer revealed from prospective clinical sequencing of 10,000 patients. Nat Med 23:703–713. doi:10.1038/nm.4333

Zhang L, Ren F, Zhang Q, Chen Y, Wang B, Jiang J. 2008. The TEAD/TEF family of transcription factor Scalloped mediates Hippo signaling in organ size control. Dev Cell 14:377–387. doi:10.1016/j.devcel.2008.01.006

Zhang W, Gao Y, Li P, Shi Z, Guo T, Li Fei, Han X, Feng Y, Zheng C, Wang Z, Li Fuming, Chen H, Zhou Z, Zhang L, Ji H. 2014. VGLL4 functions as a new tumor suppressor in lung cancer by negatively regulating the YAP-TEAD transcriptional complex. Cell Res 24:331–343. doi:10.1038/cr.2014.10

Zhao B, Li L, Lei Q, Guan K-L. 2010a. The Hippo-YAP pathway in organ size control and tumorigenesis: an updated version. Genes Dev 24:862–874. doi:10.1101/gad.1909210

Zhao B, Li L, Tumaneng K, Wang C-Y, Guan K-L. 2010b. A coordinated phosphorylation by Lats and CK1 regulates YAP stability through SCF(beta-TRCP). Genes Dev 24:72–85. doi:10.1101/gad.1843810

Zhao B, Ye X, Yu J, Li L, Li W, Li S, Yu J, Lin JD, Wang C-Y, Chinnaiyan AM, Lai Z-C, Guan K-L. 2008. TEAD mediates YAP-dependent gene induction and growth control. Genes & Development 22:1962–1971. doi:10.1101/gad.1664408

Zheng B, Zhu S, Wu X. 2015. Clickable analogue of cerulenin as chemical probe to explore protein palmitoylation. ACS Chem Biol 10:115–121. doi:10.1021/cb500758s

Zheng Y, Pan D. 2019. The Hippo Signaling Pathway in Development and Disease. Developmental Cell 50:264–282. doi:10.1016/j.devcel.2019.06.003

